# Maximum vertical height during wing flapping of laying hens captured with a depth camera

**DOI:** 10.1101/2024.10.12.617995

**Authors:** Tessa Grebey, Valentina Bongiorno, Junjie Han, Juan Steibel, Janice M. Siegford

**Author notes:** Corresponding author; (JMS). East Lansing, Michigan, United States of America (Tessa Grebey).

## Abstract

Cage-free housing systems for laying hens, and their accompanying guidelines, legislation, and audits, are becoming more common around the world. Cage-free regulations often specify requirements for floor space and cage height, but the availability of three-dimensional space can vary depending on system configurations. Little research has looked at how much vertical space a hen occupies while flapping her wings, which is arguably her most space-intensive behavior. Therefore, the objective of this study was to use a depth sensing camera to measure the maximum vertical height hens reach when wing flapping without physical obstructions. Twenty-eight individually caged Hy-line W36 hens at 45 weeks of age were evaluated. A ceiling-mounted depth camera was centered above a test pen and calibrated prior to collecting data. During testing, one hen at a time was placed in the test pen and recorded flapping her wings. From depth footage, the minimum distance between pixels was obtained for each frame, and we computed the maximum vertical height reached by each hen. Results for vertical space used during a wing flapping event showed that hens reached a maximum height of 51.0 ± 4.7 cm. No physical measures correlated with maximum height obtained from the depth camera (P>0.05). Hens in this study were from a single strain, were old enough to have keel damage, and were cage-reared and housed, preventing us from generalizing the results too far. However, depth cameras provide a useful approach to measure how much space laying hens of varying strains, ages, and rearing/housing methods need to perform dynamic behaviors.

## Introduction

Worldwide, about 3 billion of the 7.5 billion total laying hens are kept in cage-free housing systems [1]. The percentage of hens in cage-free systems varies among regions, reaching up to 99% in some European countries, and housing a substantial share of the hen population in other countries including Egypt (50%); Australia (45%); the United States (40%); Colombia, Nigeria, and Ghana (30%); South Africa (14%); and Canada (13%) [1; 2]. In some cases, cage-free systems may be implemented because of economic necessity or because they suit production needs of a region, in others the push to end the use of caged systems is driven by those advocating for improved welfare for hens on commercial farms [3; 4; 5; 6]. For example, in a survey 70% of U.S. consumers stated their support for a shift to cage-free eggs [7]. Pushes to house hens in cage-free systems for welfare reasons often result in standards, certifications, laws, and corporate commitments intended to support this objective. At the heart of many of these policies or laws, are decreased stocking density (more space per hen) and provision of resources to allow hens perform key behaviors important to welfare [3; 8; 9; 10].

In the United States, there are no federal regulations covering laying hen housing systems used by the egg industry, and the transition to cage-free eggs in the U.S. is occurring in piecemeal fashion. Several states have implemented legislation to ban conventional cages in egg production, and various corporations have made commitments to using cage-free eggs [4; 11; 12]. These state-by-state commitments often affect producers in other states or areas of the world with differing regulations who wish to import eggs to those states or for sale by those corporations. Laws passed in California, Massachusetts, Michigan, and Rhode Island, for instance, stipulate that hens must be able to fully extend their limbs without touching the side of their enclosure and eggs produced elsewhere but sold in these states must meet these criteria.

While allowing hens more freedom of movement is laudable, the lack of more precise language raises concerns for practical implementation [13]. For example, do the housing systems need to allow enough space for only a single hen to extend her limbs at a time, or must all hens in a flock be able to do so simultaneously without interference from other hens? (The latter would require far more space.) The wings of laying hens clearly require much more space to extend than their legs, particularly in the vertical plane, but little is known either about the wing dimensions of laying hens or when, why, and how often they extend them. A hen is likely fully extending her limbs not only when stretching, but also during wing flapping, which involves horizontal and vertical spreading of both wings in front of and around her body [14; 15]. Wing flapping is performed by chickens as a component of dust bathing [16], during wing-assisted jumping or running [17; 18], as part of agonistic conflict [19], during escape attempts [20], in anticipation of positive reward [21], in response to frustrated motivation [22], and on its own as a dynamic comfort behavior [14].

In 2023, the European Food Safety Authority (EFSA) Panel on Animal Health and Animal Welfare highlighted the importance of hens’ freedom of movement in cage-free systems, providing a behavioral space model to calculate the birds’ space allowance and stocking density based on specific behaviors [23]. Calculations were made with the intent to allow for unconstrained performance of behavior while reducing plumage damage, and wing flapping was used as an example [23]. Two studies that previously examined the space required by hens to wing flap were used as the basis of the EFSA wing-flapping space calculations [18; 24]. Both research groups examined the horizontal area occupied by hens while flapping, concluding that between 1693.0 ± 136.0 cm^2^ [18] and 3344.5 ± 92.3 cm^2^ [24] of floor area was needed, with brown feathered hens requiring less space than white [24]. However, only Mench and Blatchford [18] have published findings regarding the amount of vertical space used by laying hens during wing flapping, reporting Hy-Line W36 hens used 49.5 ± 1.8 cm of vertical space while flapping their wings while jumping down from a perch. However, the angle at which wings are raised, and the subsequent maximum height, varies depending on the function of the flap during locomotion (e.g., [17]). This suggests that wing flaps performed when a hen is stationary such as for the purpose of stretching, might occupy more vertical space and would be important to assess and incorporate into space calculations, guidelines, and equipment design.

Technology allows new insights into the study of animal behavior, including poultry [25; 26]. Modern imaging technology has the capacity to capture high speed motions in three dimensions and can be used to assess the amount of space occupied by a hen while wing flapping. Given that more hens are being housed cage free across the world, it is important to know how much space a hen occupies while wing flapping to assist both those developing standards or enforcing legislation as well as equipment manufacturers, builders, and producers as they design and implement aviaries. The primary objective of this study was to explore using a depth camera to measure the maximum vertical height needed for laying hens to wing flap. Secondarily, we evaluated whether physical measures of hens’ body weight and wing dimensions would correlate to the maximum wing-flapping height observed. Findings from this study may provide some idea of three-dimensional space requirements for wing flapping and provide guidance for other technology-based methods to address other space use questions in laying hens.

## Materials and methods

### Ethical approval

Prior to beginning research, study protocols were evaluated and approved by the Michigan State University Institutional Animal Care and Use Committee (PROTO202100131).

### Animals and housing

Twenty-eight Hy-Line W36 hens that were part of the fertile egg flock at the Michigan State University Poultry Teaching and Research Center in East Lansing, MI were the animals involved in this project. At the time of testing, hens were 45 weeks old. Hens were individually housed in cages that measured 30.5 x 46.5 x 42.0 cm; the floor in each cage sloped to allow eggs to be easily collected, resulting in slightly more vertical space (∼3 cm) near the front of the cages compared to the back. Prior to placement in their current cages in the fertile egg flock, the birds were reared in pullet cages.

### Physical measures from hens

Two trained persons collected physical measures of body weights and wing dimensions from each hen. Hens were removed one by one from their home cages and held upright by one person, so that their right side was pressed against the person holding them and their left wing could be handled freely by the person taking the measurement. Physical measurements were taken from the left wing of each hen. First, a ruler was held gently over the front of each hen’s folded wing to assess the length from carpal joint to the distal tip of the longest primary feather. The wing was then fully extended horizontally from the hen and the ruler was held underneath the wing to record the length of the physical structures of muscle and bone. This was done by looking up from underneath the wing so that ceiling lights shone through primary feathers to indicate where the muscle and bone ended at the distal tip of the longest phalange. Lastly, with the left wing still fully extended, a yardstick was placed against the hen’s body underneath the wing and the wing was gently flattened on top to get measurements to the distal tip of the longest primary feather. Each hen was weighed by placing her gently in a bucket on a digital scale before she was returned to her home cage.

### Depth camera and experimental set up

An Intel RealSense Depth Camera D435i (Intel Corporation, Santa Clara, CA) was used to film hens wing flapping. This depth camera (90 x 25 x 25 mm) was mounted to the ceiling in the same barn as hens’ home cages (Fig 1). A black plywood board (121 x 121 cm) was affixed to the floor directly underneath the camera; this board provided contrast against the white-feathered hens. The face of the camera was 250 cm from the surface of the plywood. Lights were placed around the testing area to prevent shadows in the camera’s field of view. Prior to collecting any data, the depth camera was calibrated to the flat surface of the plywood board using objects of known size (such as a 5-gallon bucket) to ensure the camera was correctly reporting the depth. A foldable, wire pen (122 x 122 x 122 cm) was secured around the sides of the plywood after calibrating the camera to keep hens in the camera’s field of view. The pen was open on top, giving the depth camera an unobstructed top-down view of each hen during testing, and the sides were tall enough to prevent hens from flying out. An RGB video camera (Canon VIXIA HF M41 A, Tokyo, Japan) was positioned on a tripod at hen height, facing the test pen to film each hen from a side-view as she flapped her wings. Video from this camera was used to help pinpoint when wings were extended upward during wing flapping.

**Fig 1.**
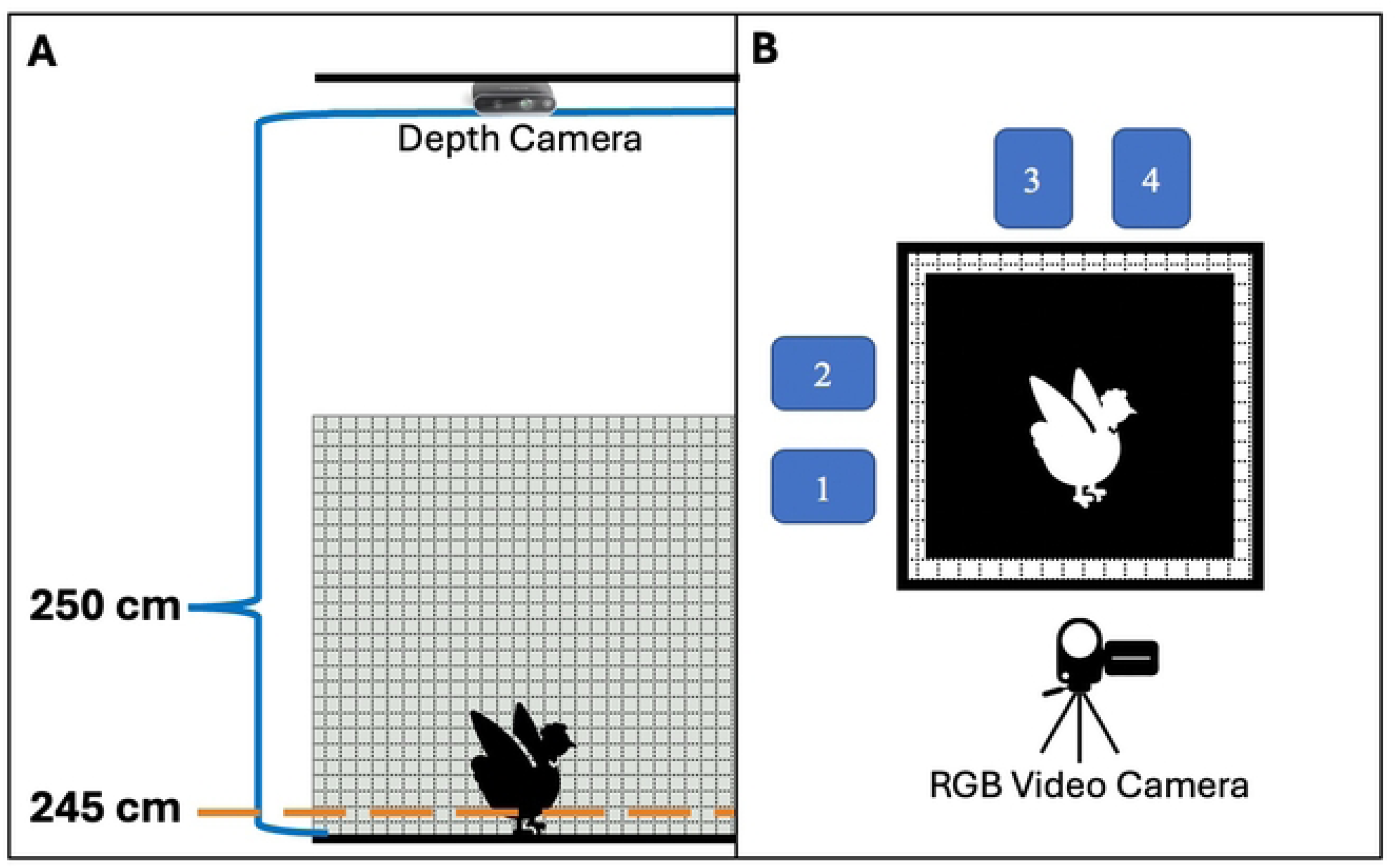
Schematics of the experimental set up. A. Side view of test pen with a hen. The depth camera was centered directly above the test pen, 250 cm above the surface of the black-painted plywood floor. The orange line represents the threshold applied for depth video analyses, which was 5 cm above the floor of the pen. B. Top-down view of the test pen. The blue rectangles labelled 1-4 represent the small carriers that hens were placed in prior to testing; these carriers faced the test pen allowing hens to see other hens during holding and testing. An RGB video camera on a tripod recorded the hen from the side to confirm wing flapping and likely time of maximum vertical extension.

We encouraged wing flapping by placing hens individually into small, hard-sided pet carriers (45.5 x 29.2 x 30.5 cm) to restrict their space prior to moving them into the pen for testing. These carriers were smaller than hens’ home cages, and we expected that rebound effect would lead to hens stretching and flapping their wings promptly once they had room to do so in the test pen. Hens spent between 10-60 minutes in the carrier, as described in more detail below in the test procedure section.

### Wing flapping experimental procedure

Two people handled hens and conducted the testing procedure. Hens were habituated to the procedure at least twice prior to data collection. All 28 hens were tested on the same day for consistency in lighting and depth calibration. Hens were removed from home cages in groups of four and were placed individually into one of the four small carriers. These carriers were situated around the test pen so that each group of four hens had visual access to one another (Fig 1).

Hens remained in the carriers for at least 10 minutes to allow study personnel to record hens’ numbers and to restrict hens to encourage quick wing flapping. After space restriction, one person removed the first hen in the group from her carrier and placed her into center of the test pen. As soon as the handler stepped away from the test pen (to avoid shadows in the image), the second person began simultaneously recording with the depth and video cameras. Each hen remained in the test pen until she flapped her wings, or 10 minutes elapsed. If a hen did not flap her wings within 10 minutes, she was returned to her carrier and video files from that attempt deleted. Once other hens in that group were tested, the non-flapping hen was placed back in the test pen for a second test attempt. When a hen successfully flapped her wings within the allotted time, the depth camera and video camera were stopped, and both video files were saved and labelled with date and hen ID. The hen was then returned to her carrier. Once all 4 hens in a group were recorded flapping their wings in the test arena, they were returned to their home cages and another group of 4 hens were removed for the test procedure. This was repeated until all 28 hens were recorded flapping their wings. No hens were kept away from their home cages (i.e., feed and water) for more than 1 hour. Between 3-7 hens were tested per hour (depending on how quickly hens flapped and whether more than one attempt was needed for a hen).

### Analyzing depth videos

Each of the 28 videos were recorded using the Intel® RealSense™ Viewer software (Intel Corporation, Santa Clara, California, USA) and saved in rosbag format. This type of file (rosbag) has a large file size compared to a typical .mp4 formatted file, but it contains all required data and meta data to re-construct an RGB image as well as a depth map. MATLAB (MathWorks, Natick, Massachusetts, USA) was utilized to read the rosbag files and to perform all subsequent processing described below. First, RBG and depth streams were aligned and frames from each video file were cropped to only feature the hen standing on the black plywood while flapping. We then performed a post-hoc analysis of depth streams and applied a distance-based threshold so that each depth video contained only those instances (pixels) when the depth values of the pixels fell in the threshold range. In this case, as the depth camera was 250 cm above the ground, the threshold was set to 245 cm. This threshold generated a binary mask to segment out pixels corresponding to the body of the hen to be included for analysis (i.e., hens’ bodies reached a high enough vertical point to be captured by the depth camera and was separated from the more distant black plywood). The binary mask was then applied to the aligned RGB frame for visual verification that no ‘hen pixels’ had been missed. Finally, all overlapped depth frames, overlapped RGB frames, and the masked RGB frames were concatenated and saved into a new .mp4 video. In the masked RGB, a red dot was placed to note the pixel that represented the highest point in each frame. In this way, when a video of a hen wing flapping is played, the red dot moves around the hen to indicate which part of her body was at the highest vertical point while wing flapping (Fig 2).

**Fig 2.**
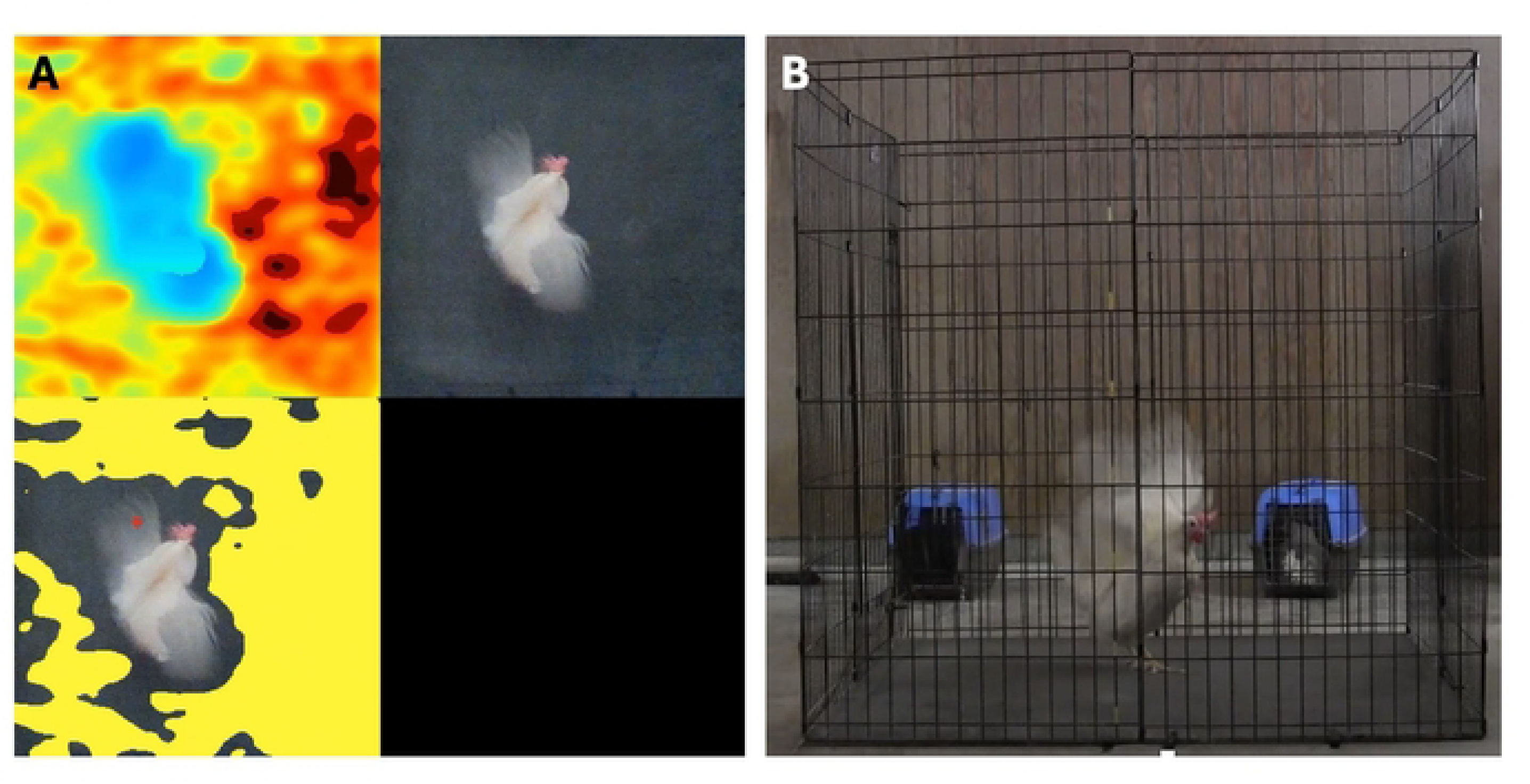
Examples of images used to measure vertical height reached by wing flapping hens while in the test pen. A. Images taken from the final depth video file. The image on the lower left panel shows the binary mask in yellow, which indicates that those pixels were not included in depth analysis. The red dot indicates the part of the hen’s body that is at the highest vertical point in each frame of the flapping event. B. Image from the RGB camera placed beside the test pen corresponding to the example images captured by the depth camera.

### Statistical analyses

The weight, length of bent wing (carpus to primary tip), extended to primary tip, and extended phalanges to phalanges were recorded from 28 hens and subjected to correlation analyses by means of IBM SPSS Statistics (International Business Machines Corporation, Armonk, New York, USA). All data were tested for normality, and as data were not normal, the non-parametric Shapiro-Wilk test was selected due to the reduced sample size. The variables were then tested for linearity, and, as the assumptions for a Pearson correlation test were not satisfied, a Spearman correlation test was performed on paired physical measures.

## Results

Once placed in the test pen, hens took between 35 seconds to over 7 minutes to successfully flap their wings. Only 2 out of the 28 hens did not flap their wings within the 10-minute time limit and had to be retested. Wing-flapping hens reached a maximum height of 51.0 ± 4.7 cm (Table 1).

**Table 1.** Vertical height reached by 28 Hy-Line W36 hens while wing flapping along with physical measures of weight and wing dimensions.

| Measurement<sup>a</sup> | Mean ± SEM (min – max) |
| --- | --- |
| Maximum vertical height during flap (cm) | $51.0 \pm 4.7$ (42.0 – 63.0) |
| Body weight (kg) | $1.6 \pm 0.9$ (1.3 – 2.1) |
| Folded wing from carpus to distal primary tip (cm) | $23.9 \pm 0.6$ (23.0 – 25.0) |
| Extended wing from body to phalanges (cm) | $16.8 \pm 1.0$ (15.0 – 19.0) |
| Extended wing from body to distal primary tip (cm) | $34.5 \pm 2.6$ (33.0 – 47.0) |

<sup>a</sup>Maximum vertical height was measured using the depth camera. Wing measurements were taken from the hen's left wing.

Averages of the various physical measurements from the 28 subject hens are presented in Table 1. No significant correlations were discovered between the maximum height obtained from the depth camera and the various physical measurements of hens (Table 2). When considering the possible combinations between the various physical measures, a marginally significant correlation was found between the length of a hen’s folded wing and the length of her extended wing extended to the tip of the longest primary feather (r=0.36, p=0.06).

**Table 2.** Spearman rank correlations between physical measurements taken from hens.

| Physical Measures Correlated | r | P-value |
| --- | --- | --- |
| Weight x Folded wing (carpus to primary tip) | 0.12 | 0.54 |
| Weight x Extended wing (to primary tip) | 0.18 | 0.38 |
| Weight x Extended wing (to phalanges) | 0.05 | 0.79 |
| Folded x Extended wing (to primary tip) | 0.36 | 0.06 |
| Folded x Extended wing (to phalanges) | 0.04 | 0.82 |
| Extended wing (to primary tip) x Extended wing (to phalanges) | 0.27 | 0.17 |
| Height<sup>a</sup> x Weight | 0.15 | 0.45 |
| Height<sup>a</sup> x Folded wing (carpus to primary tip) | -0.27 | 0.17 |
| Height<sup>a</sup> x Extended wing (to primary tip) | 0.15 | 0.46 |
| Height<sup>a</sup> x Extended wing (to phalanges) | 0.04 | 0.84 |

<sup>a</sup>Maximum height during wing flap measured using depth camera

## Discussion

The hens examined in the present study used an average of 51.0 ± 4.7 cm of vertical space while flapping their wings. Our results are similar to those reported by Mench and Blatchford [18], who found a maximum wing flapping height of 49.5 ± 1.8 cm in their Hy-Line hens. Together these findings suggest that systems offering < 50 cm of usable, vertical space between levels or between the lowest tier and the floor likely does not allow hens, at least of this strain, to wing flap without touching the ceiling in those areas.

It should be noted that our hens and those studied by Mench and Blatchford [18] were older birds (45 and ∼78 weeks of age, respectively), and had been reared and housed in cages. It is therefore probable these hens had limited or impaired mobility due to prolonged confinement and age. Moreover, it is possible that hens from our study were never afforded enough three-dimensional space to lift their wings other than the brief time they spent in the test pen, and the manner in which they flapped may not be representative of hens with more muscle development and practice wing flapping. In cage-free systems, hens have more opportunity to use and develop their muscles [27], including breast muscles that assist with wing flapping [28]. Future studies utilizing depth cameras could space use for dynamic behaviors by younger hens, especially those that were reared and housed in cage-free systems as hens with better muscle development and practice wing flapping may use more than 51 cm of vertical space.

No strong relationships were found either among the various measures of hens’ wings or between wing measures and their body weights. However, a marginally significant correlation was observed between the measurement of the folded wing and that of the extended wing (through the distal tip of the longest primary feather). A study based on a larger sample size would be needed to confirm or discard this trend toward significance. The lack of proportional relationship was surprising when comparing hens’ weight to wing measures, as wings are supportive structures in birds’ locomotion [29]. Pennycuick, [30] reported that while wing area and wingspan can vary broadly and independently from each other in several bird species, allometric proportions are typically observed between such measurements and birds’ body weight. However, we used a relatively small sample of hens with little variation in body weight and greater variation in wing measures and may not have had the power to detect a relationship. Additionally, laying hens are *Galliformes*, which are ground birds whose wing use is dedicated to bipedal movement assistance and limited short flights [29]. For this reason, in comparison with strong fliers, hens may not require a proportional variation in wing dimension in relation to their weight. As explained above, the absence of correlation between maximum vertical height while wing flapping and the birds’ weight and physical measures could also be explained by the limited vertical space available to the hens in their housing. The cage dimensions were too small to allow hens to flap their wings, certainly not with full wing extension. Such a factor may have altered their wing flapping performance in ways that do not related to underlying body weight and physical measures of their wings.

When a hen flaps her wings, she not only requires vertical space but also adequate free horizontal space, often referred to as usable floor space when discussing space requirements of cage-free hens which is critical to understand in determining stocking densities in housing systems [23; 31; 32]. Previous studies on horizontal wing flapping space use have reported hens of this same strain (Hy-Line W36) to use 3344.5 ± 92.3 cm^2^ [24], and 1693.0 ± 136.0 cm^2^ [18] of horizonal space. The variation among previous results may be explained by the rearing and housing of the hens used and conditions under which data were collected. Riddle and colleagues [24] collected measurements from hens who had been cage-free reared and housed and was done while they were performing stationary wing flaps on the floor in an open litter area of their home aviary. Mench and Blatchford [18], collected data from hens that were cage-reared and housed and was done as hens were jumping from a perch in a test chamber. Further, in Riddle et al. [24], the authors deliberately focused on the phase of the flap occupying the most horizontal area. As different wing-based behaviors require varying amounts of three-dimensional space, it is possible that hens use less room when flapping their wings for jumping purposes [18] as opposed to when they are standing stationery and flapping, when they may be more likely to fully extend their wings as part of a stretch or display [14; 19].

The amount of space a hen occupies when performing dynamic behavior also varies depending on genetic strain [24]. There are also notable genetic differences in hens’ skeletal and musculature structure that may influence how they perform specific behaviors, like wing flapping. White-feathered hens tend to have heavier breast muscles and larger keel bones, whereas brown-feathered hens have heavier leg muscles [33; 34]. More research must be done to assess space requirements across multiple strains of laying hens before comprehensive guidelines can be provided, and it is possible that a range of height guidelines are necessary for hens of different strains.

Finally, 26 of the 28 hens tested wing flapped between 35 seconds and 7 minutes. The two birds who did not wing flap during the initial 10-minute testing period did so when placed in the test arena a second time. This finding shed light on two important aspects. On one hand, the hens in this study were reared in cages, with no opportunities to exercise their wings over time. Nevertheless, in a relatively short amount of time all the birds wing flapped. This finding underlies the innate motivation for such a behavior in the hens, and the rebound need to “stretch” their wings soon after being placed in the test arena was probably consequential to cage housing. Similarly, space availability is a relevant factor shown to affect the behavioral performance of hens in previous studies [31]. On the other hand, our hens never occupied as much space while wing flapping as birds in Riddle et al. [24], despite having plenty of room to do so during the test. This observation reinforces the hypothesis that long-lasting effects of cage rearing systems may impair the birds’ physical ability to wing flap. Anecdotally, the tested hens appeared inexperienced while wing flapping; showing uncoordinated movements like slipping, sliding, and tilting that are usually observed in young pullets (personal observation).

## Additional considerations

It remains unclear if spacing requirements in cage-free legislation should require that a hen (or all hens in a group) be able to flap her wings at all times. Hens may not need or want to flap their wings at certain times of day because they are performing other behaviors that cannot occur concurrently, such as eating or laying eggs. Additionally, laying hens do not distribute themselves evenly throughout a given area but often cluster to access resources or to maintain proximity to each other [35]. When hens congregate within a housing system, they effectively increase the stocking density of that immediate area, which can reduce their freedom of movement in that space [32; 36]. Crowding may limit a hens’ actual ability to wing flap at a given time in a particular place or she may not think she is able to flap due to perceiving the proximity of her flock mates [37]. The function, motivation, and causation of wing flapping should be further examined so that cage-free initiatives can clearly drive practical improvements to cage-free husbandry and housing that genuinely benefit hens.

## Conclusion

Multiple conclusions can be drawn from the present research. For a start, depth cameras, such as the one used here, appear to be useful as a supporting technology for behavioral studies of hens. Their use allowed us to successfully capture wing flapping events and generate measurements on the space required for hens to wing flap. The space occupied by the hens while wing flapping provided interesting perspective regarding current behavioral opportunities (or lack thereof) within cage-free aviary systems, where heights between levels may be less than what a hen needs (or perceives she needs) to wing flap. Spaces beneath and within the tiers can host a transiting or standing hen but may not be spacious enough to allow comfortable expression of wing flapping behavior, therefore limiting laying hens’ welfare in the very cage-free systems intended to improve it. Finally, depth camera data and physical measures collected from the hens were mostly uncorrelated, which hampers the use of easier-to-collect physical measures from hens of different ages and strains to predict space needed for wing flapping. Further research on hens of different genetics and ages is required, particularly encompassing strains which likely display greater muscle strength and flapping abilities (e.g., local breeds and genotypes less selected for commercial egg laying).

## Acknowledgements

The authors thank the Michigan State University Poultry Teaching and Research Center manager Angelo Napolitano for his assistance with study set up and research assistant Elizabeth Gregas for her help in data collection.

## Supporting Information

**S1 File. Wing flapping depth data and physical hen measures.** Raw data analyzed and presented in the manuscript are available in the two worksheets of the supplemental file.

